# Towards robust and generalizable super-resolution generative adversarial networks for magnetic resonance neuroimaging: a cross-population approach

**DOI:** 10.1101/2022.06.13.495858

**Authors:** Leona Charlotte Förster, Lucas da Costa Campos, Martin Kocher, Svenja Caspers

## Abstract

Magnetic resonance imaging (MRI) is fundamental to neuroscience, where detailed structural brain scans improve clinical diagnoses and provide accurate neuroanatomical information. Apart from time-consuming scanning protocols, higher image resolution can be obtained with super resolution algorithms. We investigated the generalization abilities of Super Resolution Generative Adversarial Neural Networks (SRGANs) across different populations. T1-weighted scans from three large cohorts were used, spanning older subjects, newborns, and patients with brain tumor- or treatment-induced tissue changes. Upsampling quality was validated using synthetic and anatomical metrics. Models were first trained on each cohort, yielding high image quality and anatomical fidelity. When applied across cohorts, no artifacts were introduced by the SRGANs. SRGANs that were trained on a dataset combining all cohorts also did not induce any population-based artifacts. We showed that SRGANs provide a prime example of robust AI, where application on unseen populations did not introduce artifacts due to training data bias (e.g., insertion or removal of tumor-related signals and contrast inversion). This is an important step in the deployment of SRGANs in real-world settings.

## 1. Introduction

One of the fundamental tradeoffs in magnetic resonance imaging (MRI) is the balance between image resolution, acquisition time, and signal-to-noise ratio^1^. High resolution images are naturally more desirable as they provide more detailed information for use in brain research and diagnostics. However, in order to obtain them, long-lasting data acquisition procedures are typically necessary, which can be uncomfortable for patients and comes at either higher operational costs or a decrease in signal-to-noise ratio^2^. Participants’ unwillingness to take part in long imaging sessions can also lead to damaged or incomplete scans. Upsampling, a method for increasing data resolution during post processing, has been proposed as a way out of this dilemma^1^. However, a biased upsampling mechanism carries the risk of attempting to restore patterns that were never present in the original images, resulting in misclassifications in downstream analyses. For example, brain tumors can cause unpredictable alterations in the MR signal of the surrounding tissue, and neonatal brains are characterized by an opposite MR contrast compared to adult brains, with gray matter regions creating stronger MR signals than the white matter due to incomplete myelination. The risk for a biased upsampling algorithm to unintentionally add or remove these features hinders the deployment of a universally applicable upsampling method.

The history of image-upsampling methods is characterized by three methodological milestones. Firstly, interpolation-based methods have been used since the inception of magnetic-resonance imaging^3^, where cubic spline interpolation in particular found widespread adoption due to its simplicity and speed^3^. However, its inability to adapt to domain-specific optimal strategies limits its capacity to recover delicate structures. Interpolation methods were superseded by the second milestone, namely patch-based (also often called dictionary-based) algorithms which extract details contained in the low-frequency components of the image in order to reconstruct high-frequency information^4–6^. However, ad-hoc adaptations of these algorithms had to be developed for each application. The third milestone is the usage of Deep Neural Networks (DNNs), which has led to major breakthroughs in the field. DNNs have led to drastic improvements in many areas^7^, especially in computer vision tasks which are particularly amenable to Convolutional Neural Networks (CNNs)^8^ which have found widespread usage in classification^9^, segmentation^10^, and image generation^11–13^.

The usage of Super resolution CNNs, an image upsampling architecture^14,15^, for MRI scans has recently been validated^16^, leading the way to many studies on the application of Deep CNNs for this purpose^17–20^. A further leap forward in image upsampling was achieved by combining CNNs with a discriminator network, in what is known as the Generative Adversarial Network (GAN) framework^21^. Originally developed as a tool for the generation of new scenes and faces, GANs are able to learn the complex underlying feature-space of real data through adversarial training and can create true-to-life images. This makes GANs an especially useful framework for medical imaging^22^, where they have found application in denoising^23^, reconstruction^24^, modality transfer (e.g., generating T1w MR images to and from T2w MR images^25^ or CT scans from MR data^26^, among many others.

Despite these advances, the application of GANs to MRI upsampling is still at its infancy. Refs.^27,28^ provided early indications that GAN-based upsampling can lead to state-of-the-art performance in image upsampling as indicated by image metrics. Nonetheless, a fundamental prerequisite of training a Machine Learning (ML) model is that the training population needs to be tailored to the problem at hand lest biases in training data can lead to incorrect results. This is particularly important in the case of GANs, which are prone to hallucinating features in and out of images^21,29,30^. If GAN-based upsampling is to be used in clinical settings, it needs to be validated against population-induced artifacts. In fact, Ref.^31^ provides a harrowing tale of GANs removing tumors from patients’ scans, which would be disastrous in a clinical setting.

In this work, we trained 3D Super Resolution GANs (SRGANs)^27,30^ and studied their resiliency against population-based artifacts, with special focus on anatomical metrics and features, e.g., MR contrast inversions or even the removal/insertion of tumor- or treatment-induced tissue changes. The abilities of SRGANs to handle varied image types were probed, as well as their usability in clinical and scientific settings. In order to cover a wide array of clinically relevant anatomical features and common use cases for MRI data, T1-weighted scans were used. We aimed at including datasets with a high degree of diversity in population age, brain health, and morphology. To include samples with such different base characteristics and thus, potential sources of bias, data from a large healthy population-based imaging dataset (1000BRAINS), the Developing Human Connectome Project (dHCP) that hosts images of neonatal brains, and large sample of datasets from patients with brain tumors (gliomas) were used. SRGANs were trained to perform upsampling of MRI scans with shape 80 × 98 × 84, doubling its linear resolution to 160 × 196 × 168. The upsampled quality was evaluated using both image quality metrics (peak signal to noise ratio (PSNR) and structural similarity index (SSIM)) and domain-specific metrics (cortical thickness, cortical surface area and brain volume). Domain-specific metrics were calculated using the FastSurfer software package^32^. The SRGANs trained in this work were also made available in an easy-to-use and open-source software package.

## 2. Results

### 2.1. Experiment 1: Super Resolution within the 1000BRAINS cohort

In this experiment, we train SRGANs with an increasing number of subjects from the 1000BRAINS training dataset (for methodological details see Sec 5.6). When using only 10% of the original dataset size, the resulting PSNR ranged between 22.87 and 24.71dB, (24.12% to 34.07% increase compared to the interpolation-based method; see Table 2 for baseline) and SSIM ranged from 0.83 to 0.86 (3.05% to 7.71% increase) (see Fig. 1). Four of the 16 SRGANs showed abnormally low performance (PSNR < 24dB). These underperforming networks were all trained on the same subsampling, revealing that with 10% of the data (99 participants) there is a real possibility of an unrepresentative or incomplete sampling of the population, resulting in subpar upsampling quality. No large changes in the average PSNR and SSIM were observed as the number of subjects was increased from 99 all the way up to 985. However, the gap between the best and worst performing networks decreased noticeably. One remarkable exception is that a single network trained on a subsampling with 247 subjects reached a maximum PSNR of 26.60dB. This, however, was not indicative of a more general tendency.

**Table 1:** Comparison of anatomical measurements for training sessions with 985 subjects in Experiment 1 and with 1495 subjects in Experiment 4 (see Sec. 2.4). Asterisks indicate the level of statistical significance (ns: p > 0.05, *: p ≤ 0.05, **: p ≤ 0.01, ***: p ≤ 0.001). All values are given as mean ± std.

|  | Diff to HighRes (%) |
| --- | --- |
| **Tricubic** |  |
| Cortical Thick. (LH) | -18.71±2.58 *** |
| Cortical Thick. (RH) | -19.10±3.12 *** |
| Cortical Area (LH) | -1.62±3.02 *** |
| Cortical Area (RH) | -3.55±3.79 *** |
| Brain Volume | 4.22±5.06 *** |
| **SRGANs (1000BRAINS; Exp. 1)** |  |
| Cortical Thick. (LH) | -18.94±2.01 *** |
| Cortical Thick. (RH) | -18.18±2.33 *** |
| Cortical Area (LH) | 2.29±3.71 *** |
| Cortical Area (RH) | -0.65±3.95 ** |
| Brain Volume | 0.95±5.17 * |
| **SRGAN (Generalist; Exp. 4)** |  |
| Cortical Thick. (LH) | -16.12±2.02 *** |
| Cortical Thick. (RH) | -16.16±2.29 *** |
| Cortical Area (LH) | 3.12±3.17 *** |
| Cortical Area (RH) | 0.95±3.59 ** |
| Brain Volume | 2.25±2.49 *** |

**Table 2:** Image quality of upsampled images for each cohort in Experiments 1 and 2. For Experiment 1, only the data for training with the full sample is shown. All values are given as mean ± std. n = 247.

| Cohort | Tricubic | SRGANs | Improvement |
| --- | --- | --- | --- |
| **1000BRAINS**<br>PNSR (dB)<br>SSIM | 18.44±4.15<br>0.80±0.08 | 23.86±0.05<br>0.85±0.00 | 5.42 (29.43%)<br>0.05 (05.62%) |
| **dHCP**<br>PNSR (dB)<br>SSIM | 14.15±1.64<br>0.68±0.03 | 23.03±0.08<br>0.80±0.00 | 8.88 (62.68%)<br>0.12 (18.54%) |
| **Brain Tumor**<br>PNSR (dB)<br>SSIM | 15.58±2.61<br>0.77±0.08 | 23.19±0.34<br>0.84±0.01 | 7.61 (49.81%)<br>0.07 (09.95%) |

**Figure 1:**
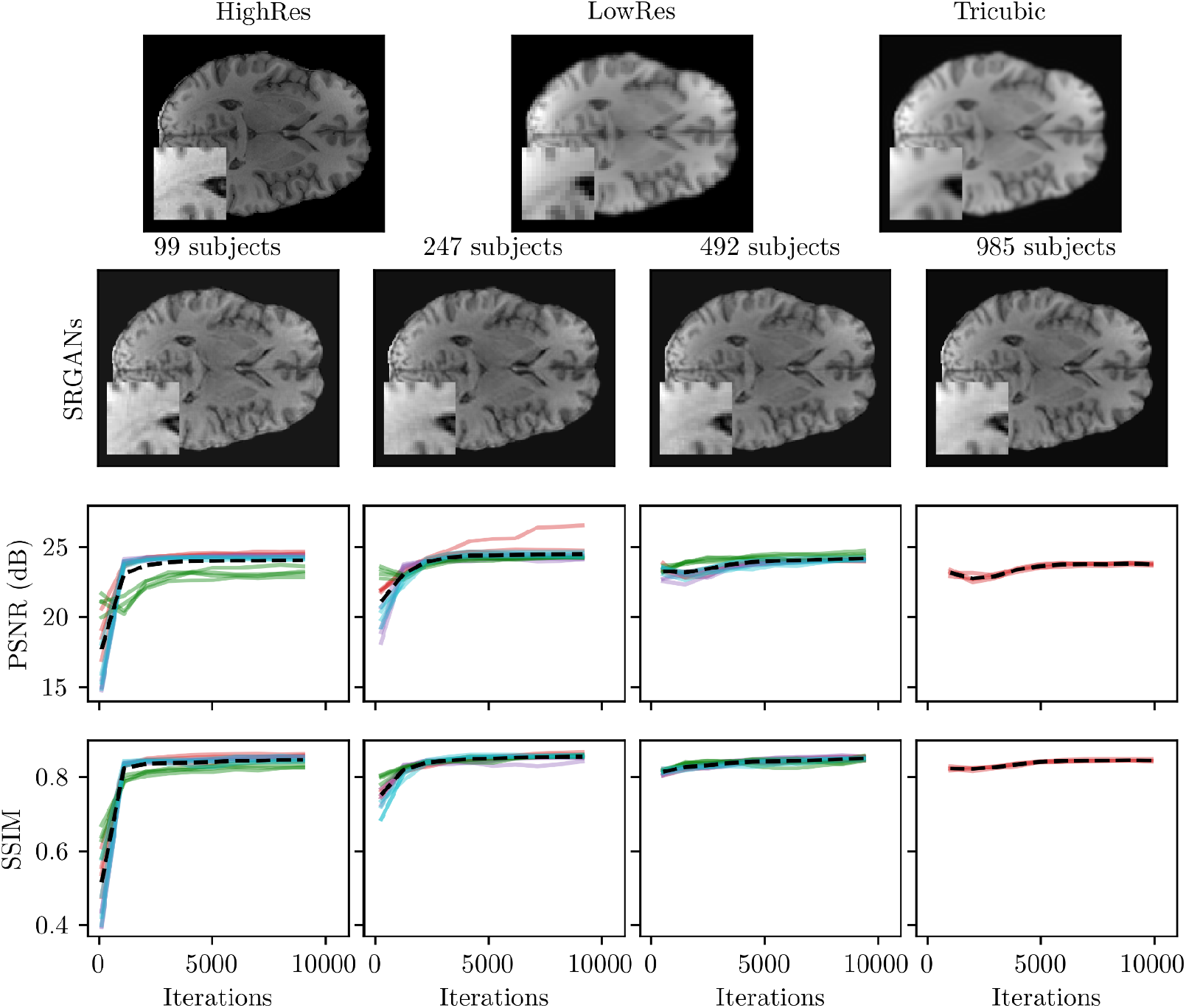
Anatomical and Synthetic metrics of varying number of data points. Each color represents a different sampling encompassing four tries with different starting weights. [Full width]

We also examined the effect of dataset-size on differences in anatomical metrics. Anatomical metrics were calculated for scans upsampled both by SRGANs and by tricubic spline interpolation. These scores were then compared with anatomical metrics calculated using the original image as baseline. Similarly to the synthetic metrics, the resulting measurements did not depend strongly on the size of the training dataset (see Supplementary Information, Table S2).

For every sample size we observed a large deviation between the cortical thickness measured from the upsampled image *I_SR_* and that from the original image *I_HR_*. For both tricubic spline and SRGAN, the downsampling and subsequent upsampling led to a loss of approximately 19% of cortical thickness (see Table 1). Additionally, the difference between the performance of the spline-based upsampling and the SRGAN-based approach were small (< 1.5%) and frequently only significant for one hemisphere. Cortical surface area presented significant but limited differences between the upsampled and original high resolution images, consisting of either small increases in the measured surface area (≈ 2%), or a slight decrease (< 1%). Unlike with cortical thickness, the comparison between the interpolation-based upsampling and the SRGAN-based method was highly statistically significant. Of the three anatomical metrics, total brain volume showed the largest difference between the two methods. Whereas the difference in measured brain volume between the SRGAN approach and the high quality image was below 1%, measurements of the tricubic approach and high quality image differed by about 4%.

### 2.2. Experiment 2: Super Resolution within the Brain Tumor and dHCP cohorts

SRGANs were trained and evaluated on the two remaining cohorts, Brain Tumor and dHCP. The full training samples as shown in Table S1 were used. These results were compared with those from SRGANs from Experiment 1 trained on the whole 1000BRAINS training dataset.

Both dHCP and Brain Tumor cohorts were amenable to SRGAN-based upsampling, reaching PSNRs in the range 23.03 to 23.86 dB and SSIMs in the range 0.80 to 0.84 (see Fig. 2). Most noticeably, the final values of PSNR and SSIM remained remarkably stable across all three tested cohorts compared to the Tricubic interpolation values, which ranged between 14.14-18.44 dB for PSNR and 0.68-0.80 for SSIM. The SRGAN method showed a much larger degree of improvement compared to the interpolation-based method. By using SRGANs, an improvement of 29% was achieved for the 1000BRAINS cohort, which showed the largest PSNR but smallest relative improvement. The other two datasets exhibited PSNR improvements of 63% and 50% for the dHCP and Brain Tumor cohorts, respectively.

**Figure 2:**
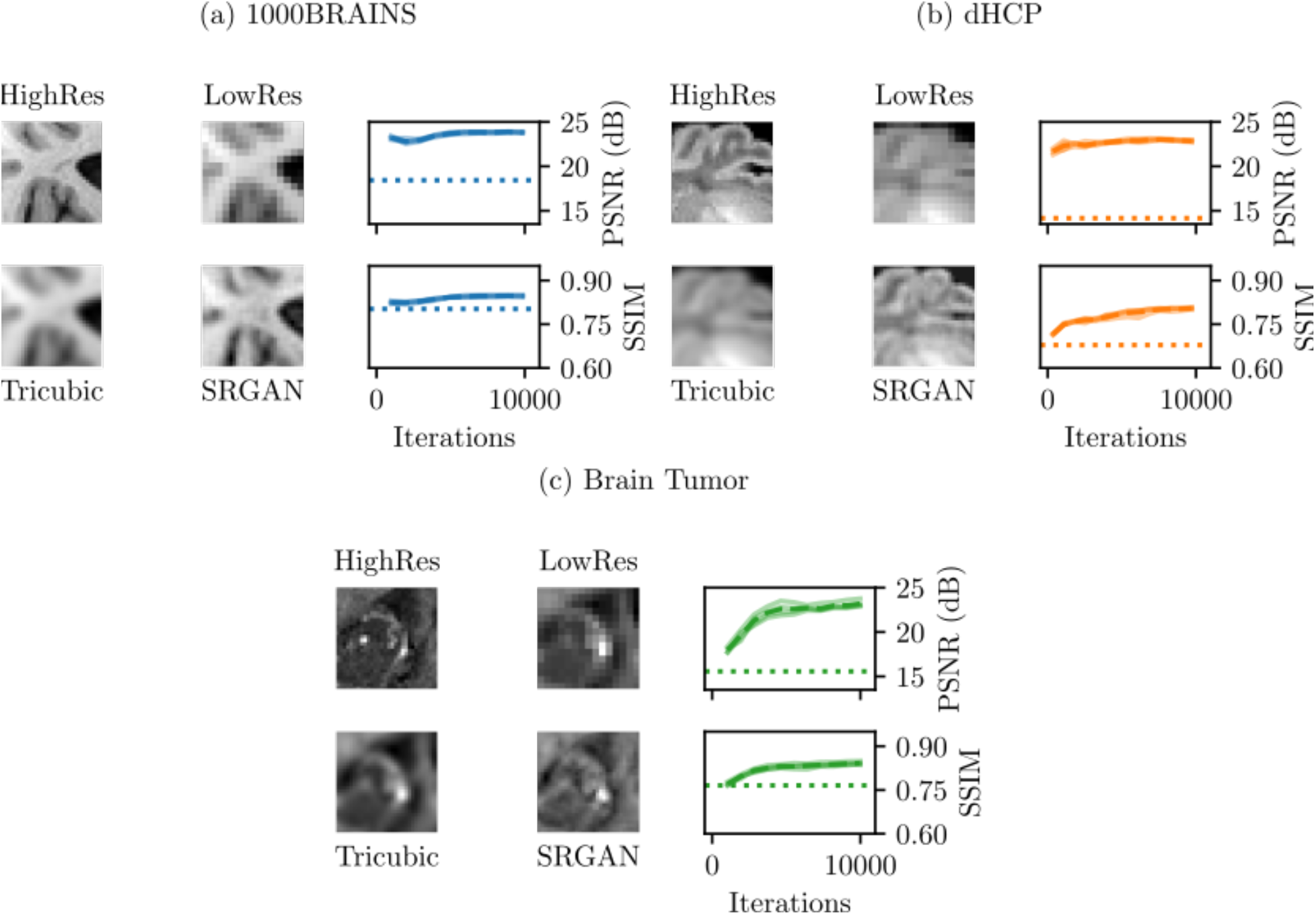
Input and output image details (of the original high-resolution image and image upsampled via SRGANs) and synthetic metrics for each cohort, as indicated within the figure. Solid lines represent independent training sessions, while broken lines depict overall averages and dotted lines show the results from tricubic interpolation. For easier comparison, some results Fig. 1 are replicated here. [Full width]

The SSIM told a similar story, with SRGAN-based upsampling of 1000BRAINS showing the least improvement (< 6%) compared with the tricubic interpolation, while the same process applied to dHCP and Brain Tumor led to SSIM improvements of 10% and 19% respectively (see Table 2).

### 2.3. Experiment 3: Super Resolution across populations

The SRGANs trained in Experiments 1 and 2 were now used to perform upsampling across datasets (e.g., SRGANs trained on the 1000BRAINs dataset were used to perform upsampling of the dHCP dataset).

The quality of the resulting SRGAN-generated images was not perceptually different from that obtained during the self-trainings (See Fig. 3 and compare with Fig. 2). Synthetic metrics indicated small differences in image quality. Interestingly, often the best image quality was not obtained by the SRGANs trained and tested on the same cohort (see Table 3). For instance, the best results when upsampling 1000BRAINS data (PSNR: 24.14 dB/SSIM: 0.85) were obtained by using the SRGANs trained exclusively on the Brain Tumor dataset. Similarly, the best PSNR on the Brain

**Figure 3:**
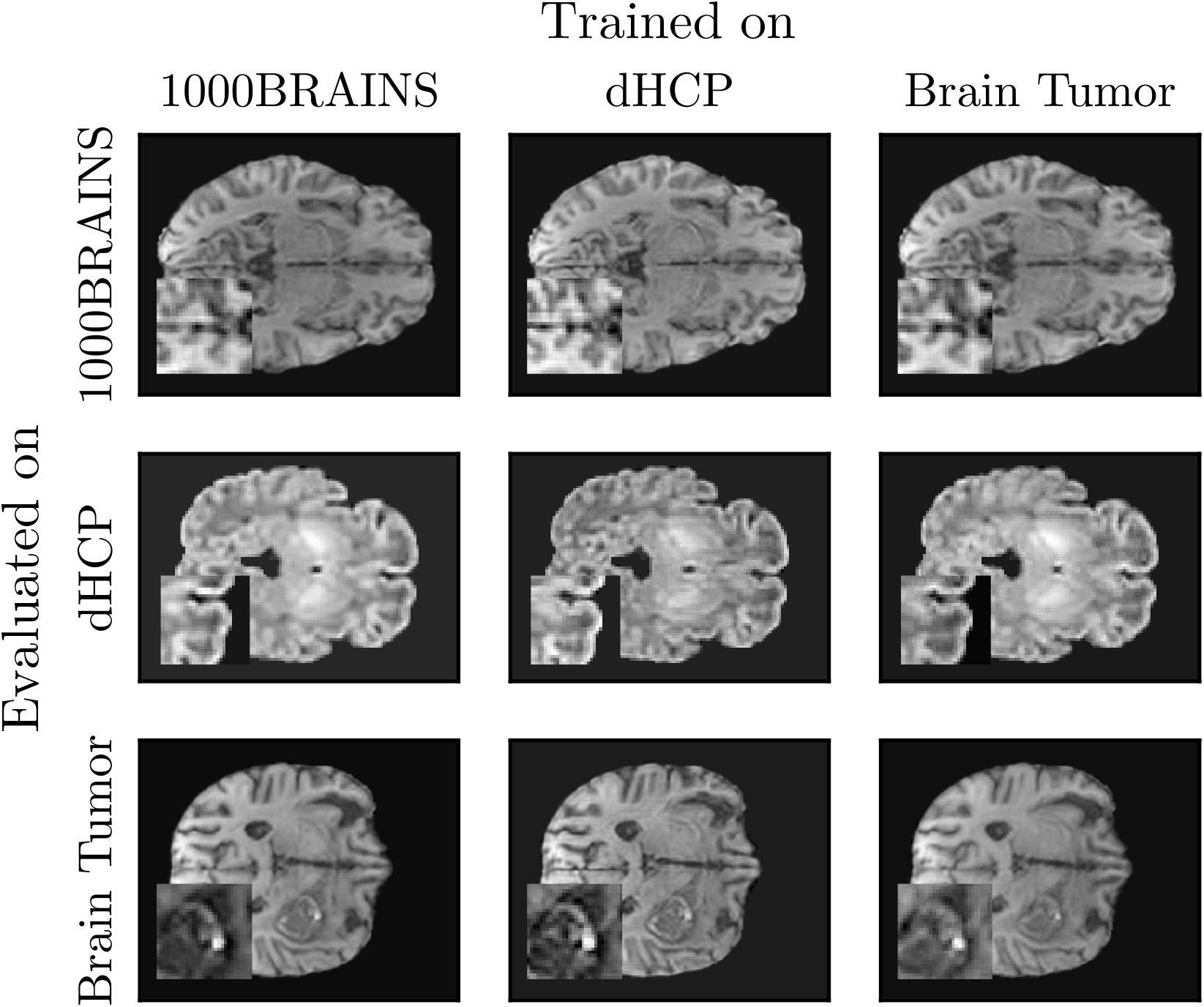
Examples of upsampled images after cross evaluation. [One-column-and-half width]

**Table 3:**
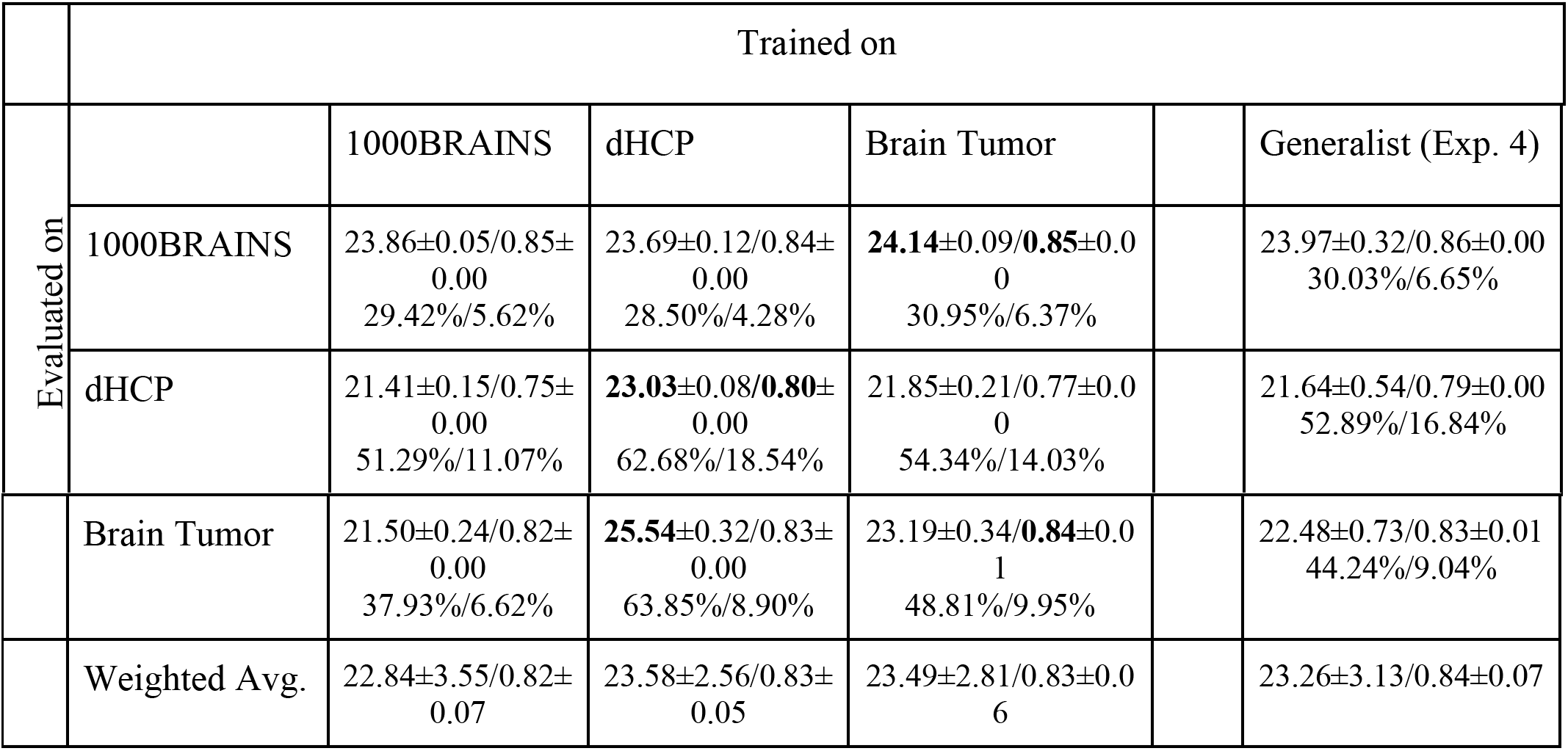
Synthetic benchmarks of Experiments 3 and 4 (see Sec. 2.4). The highest score for PNSR/SSIM is shown. Each column indicates the dataset on which the networks were trained, and each row indicates the dataset on which they were evaluated. Within each cell the absolute scores (above) and improvement relative to tricubic interpolation are shown (below, as percentage; see Table 2 for the baseline). The highest score for each evaluation dataset is written in bold. The weighted average performance of each network is shown in the last row. Values are shown as mean ± std. For sample sizes, see Table S1.

Tumor cohort was obtained from a network trained with dHCP. Overall, the networks trained on the dHCP dataset showed the largest improvements, closely followed by those trained on the Brain Tumor dataset. The SRGANs trained on 1000BRAINS yielded the smallest increase in image quality. Visual inspection of the SRGAN-generated images revealed that no insertions of tumor-related image changes into healthy adult brains or contrast inversions (as potentially possible due to the inverted myelin contrast in the infant brains of the dHCP cohort) were performed by the SRGANs. Complementarily, all networks preserved tumor-related image changes as well as the characteristic contrast of neonatal brains, even when those structures were not part of their training data.

### 2.4. Experiment 4: Super Resolution with mixed Populations

We evaluated the benefits of training a generalist network on a large dataset consisting of all scans dedicated to training, i.e., 985 1000BRAINS scans, 360 dHCP scans, and 149 Brain Tumor scans for a total of 1494 scans. Images upsampled by SRGANs trained on this combined dataset showed similar PSNRs as those in Experiment 3, hovering between 21.64 and 23.97 dB, with an average score of 23.26 dB (Table 3, last column). This was a lower average PSNR than the one obtained with the dHCP-trained network. However, the generalist network achieved the best SSIM score, ranging between 0.79 and 0.86, with weighted average of 0.84. According to the same metric, the generalist network was also the best performing SRGAN for the 1000BRAINS data, with a SSIM of 0.86 (see Table 3).

In addition, anatomical measurements were taken for the 1000BRAINS scans upsampled by the generalist cohort. There was a decrease of 16% in the measured cortical thickness in the SRGANs-upsampled images compared to the same measurement in the original images (see Table 1). This indicated a ~3% improvement compared to the images which were upsampled using the SRGANs trained in Experiment 1 and thus on 1000BRAINS alone. Cortical surface area followed the opposite trend, being overestimated by between 1% and 3% compared with the original images, a minor downgrade from the [-0.65%, 2.29%] range found in Experiment 1. This was mirrored in the brain volume, which was overestimated by more than 2% for the generalist networks, and less than 1% in Experiment 1. However, most meaningful for this work, these generalist SRGANs also did not induce any population-based artifacts, such as additional tumor-related signal, removal of tumor-related image changes, or contrast inversion.

## 3. Software and SRGAN release

Brainhance, a software package for MRI upsampling using the SRGANs trained in Sec. 2, is made available as an open source software^33^. Brainhance has been uploaded to PyPi, facilitating the installation in a local Python environment using *pip install brainhance*. The best performing SRGANs for each training dataset are released together with this software and enable upsampling of data in the absence of adequate training data. They are accessible through command line flags. Brainhance is written using the TensorFlow framework^34^, which allows the efficient usage of GPUs or CPUs. It can be used for single images, allowing for a simple user experience, however it also provides a batch mode, enabling the user to feed a list of input files into the SRGAN. This mode allows the SRGAN to be loaded into memory only once, speeding up the execution of any upsampling after the first one. Our tests indicated that upsampling takes about 1s per scan in batch mode using a single Nvidia P100 and roughly 4s per scan when using 2×Intel Xeon Gold 5120.

## 4. Discussion

We have analyzed the effects of bias on super resolution GANs using datasets drawn from three different populations: Neonatal (dHCP), older adults (1000BRAINS), and patients with brain tumor- or treatment-induced tissue changes. This work gave further empirical evidence on the reliability of Deep Learning-based upsampling in neuroimaging, especially in the context of generative models. As far as the authors are aware, no other work has analyzed the cross population safety of SRGAN-based MRI upsampling. We found SRGANs to be robust against changes within and across populations. This is particularly important in light of the work by Cohen et al.^31^. In this work, a generative model called CycleGANs was trained to perform cross-modality conversion between FLAIR and T1w data, and vice-versa. It was observed that this model was prone to the removal of tumor-related signal changes from the images, which provides a strong warning against using generated images in a clinical setting. This weakness was not observed in the current work, where the SRGANs model proved to be remarkably stable, with no tumor- or treatment-induced tissue changes removal or addition and no MR contrast issues being observed in any of the trained networks.

Initially, we examined the impact of different dataset sizes on synthetic and anatomical metrics for the 1000BRAINS cohort. In general, we observed an increase of up to 40% in PSNR and 8% in SSIM as compared to a typical upsampling method such as tricubic interpolation, in line with the findings of previous works in the field^16,27^. Measurements of brain volume and cortical area presented mixed results, with small (in the range ±3%) but statistically significant differences between the upsampled and original images. The recovered images did not, however, capture the stark signal gradient between the gray and white matter, leading to measurements largely underestimating the cortical thickness of the subject. Further methodological advancements of SRGANs are necessary to tackle this issue.

We observed a rapid saturation in all metrics with increasing sample sizes, indicating that for the purposes of MRI super resolution, a dataset of size smaller than 100 subjects already yields acceptable results in most cases. This lends further strength to a large body of previous literature that used training populations with sizes between several dozens to the low hundreds subjects^35–39^.

Subsequently, networks were trained and evaluated on the remaining two cohorts, where it was observed that the image quality improvements were cohort-dependent, with dHCP data seeing the largest improvement compared to the baseline situation of the tricubic approach. These results show that SRGANs provide a similar degree of image quality even in the presence of atypical brain scans. The previously trained networks were then used for cross evaluation (i.e., using networks trained on a given dataset to upsample data from a different cohort). Despite the population differences between the datasets, all three of them contained sufficient information to convey underlying image-feature distributions of the other cohorts. Most importantly, this experiment showed that the performance of SRGANs is robust against changes in the local dataset, and that no large, clinically relevant artifacts were hallucinated into images.

Finally, training was performed using all datasets simultaneously. Results showed similar increases in the synthetic benchmark and no population-based artifacts, as before. To the best of the authors’ knowledge, this is the largest MRI upsampling population to date (narrowly larger than Ref.^40^, where 1520 subjects were used in total). Comparing the generalist’s anatomical measurements with the anatomical scores of networks trained solely on 1000BRAINS showed slightly better cortical thickness estimates (−16%, compared to −18% with only 1000BRAINS), but higher overestimation of total brain volume (2.25% compared to 0.95%).

Most studies take a “general view” of the images, using either a single cohort or groups of comparable cohorts^18,20^, most often using synthetic metrics, such as PSNR or SSIM. As shown here, these metrics do not tell the whole story, in particular regarding domain-specific (in this case, anatomical) fidelity. The results of this work highlight that improvements in image quality metrics do not necessarily translate into more precise anatomical measurements. A similar point has been made in Ref.^22^ in regards to image quality. Therein, the authors make the recommendation that downstream measures should be used as a more reliable validation step for the generated sample. Our results further emphasize this advice.

Most importantly, this work provides an important step in the deployment of SRGANs for clinical uses by showing that their usage can be appropriate for tumor diagnosis, in so far as it does not significantly modify the shape or size of the tumor- or treatment-induced tissue changes in the upsampled image. However, due to deviations observed in anatomical measurements - particularly cortical thickness - our findings also highlight that usage of further upsampling analyses needs to be carefully considered, and that subsequent studies should be benchmarked against their specific intended usage.

Our results might also benefit population-based and clinical cohort imaging where guaranteeing stable image quality and resolution across subjects can be challenging. We expect the SRGANs stability found here to be also useful beyond the realm of neuroscience. Some particularly interesting applications are those where high temporal resolution is necessary and which therefore can be extremely limited in terms of signal-to-noise ratios. For instance, perfusion MRI is sensitive to the local microvasculature which makes the characterization of several diseases possible^41^. However, this technique requires fast scans in order to correctly estimate the passage of the contrasting agent. Another interesting possible beneficiary of stable upsampling techniques is cardiac MRI which also requires high temporal resolution to resolve the beating motion of the heart^42^.

In the current work, we studied a single super-resolution scheme (i.e., SRGANs using resize convolution). Different schemes undoubtedly present different advantages and disadvantages. While we expected the results in the current work to be extensible to other CNN-based upsampling schemes (for instance, using variational autoencoders) that take pixel-by-pixel comparisons in their losses, analyses on other methods still have to be carried out. Only T1-weighted data was used for this work, but different MRI modalities are able to capture different brain features. For instance, T2-weighted images highlight water content, as opposed to T1-weighted images, which highlight fat content; diffusion MRI captures the water diffusivity in the brain and indirectly the axonal fiber distribution. A similar study combining features from different MRI modalities would therefore be desirable.

## 5. Methods

### 5.1. Datasets

Three datasets were selected and are summarized below.

- Brain Tumor^43^: Cohort composed of patients with malignant glioma as well as treatment-induced changes, aged 28 to 81. n = 187 scans were used.
- Developing Human Connectome Project (dHCP)^44^: Cohort of neonatal subjects, with conceptional ages between 29 and 45 weeks. n = 451 scans were used.
- 1000BRAINS^45^: Population-based cohort, mostly from older participants. n = 1233 scans were used, from subjects with ages between 18 and 85 years.

### 5.2. Preprocessing Steps

The data from the 1000BRAINS and Brain Tumor cohorts were normalized using the following procedure: Initially, each scan was converted to neurological orientation if necessary. Images were resliced to size 1mm × 1mm × 1mm with dipy 1.0.0^46^ which was necessary for the dHCP dataset, as they have double the resolution of the other cohorts and to correct any minor anisotropy found in the Brain Tumor cohorts. Brain extraction masks were then generated using hd-bet version 1.0^47^.

The excess background resulting from skull extraction was cropped, and the scans were padded to a standard 160 × 196 × 168 shape using MRtrix’s 0.3.15-32 mrcrop command^48^. The resulting images were normalized in the range [0, 1]. Following Ref.^16^, the low resolution versions (LR) of the original images were generated by first performing a Gaussian blur (σ = 1) and then using the zoom function as implemented in scipy v1.5.0^49^ for downsampling by a factor two in each direction. Tricubic interpolation was performed to obtain the baseline upsampling method the networks were compared against. The entire preprocessing pipeline is summarized in Supplementary Fig. 1 (a). Due to the intensity inversion from lack of myelination in the neonatal brain, hd-bet fails to process dHCP T1w scans. Instead, the masks available as part of the dHCP data release were used (see Supplementary Fig. 1 (b)).

Following standard procedure, each of the three datasets was split into two disjoint subsets: a larger set comprising 80% of the data, which was used for training, and a smaller set comprising the remaining 20% of the data, which was reserved for evaluating the quality of generated images (see Table S1). Due to memory restrictions, the downsampled images were split into overlapping octants, each with shape 56 × 65 × 58 (112 × 130 × 116 for original resolution images) before being fed into the neural networks.

### 5.3. Network Architecture

We used SRGANs to perform supersampling of MRI samples. The 2D-basis for the model was provided by Ref.^30^ and adapted by Ref.^27^ to make it applicable for 3D MRI data. The system was composed of two distinct neural networks, a generator and a discriminator^21^. The generator was optimized to imitate the real, high resolution, data distribution (HR), i.e., to provide a realistic super resolution output image (SR) for low resolution input (LR). The discriminator was optimized to classify scans as either real data or an imitation created by the generator. In every training iteration the generator’s error was measured by its ability to create imitations the discriminator was not able to identify. The resulting learning process is commonly referred to as adversarial learning.

#### 5.3.1. Generator architecture

The generator used 3D up-convolution to create super-resolution (Suppl. Figure 2). First, the low-resolution volumes were fed into the initial 3D convolution and batch-normalization layer, followed by 4 residual blocks. Each residual block contains two 3D convolution layers, with 3 × 3 × 3 kernels, strides of 1 in every direction, and 32 output feature-maps. Subsequently, batch normalization was performed in every block to improve the network’s stability during training. The element wise sum operation at the end of each residual block combines the block’s output with unaltered information from previous layers to enable the network to learn features on all abstraction levels^30^. After the four complete residual blocks which were constructed based on the aforementioned scheme, another, differently designed residual block consisting of 3D convolution, batch-normalization and an element-wise sum component was integrated into the model. The Leaky Rectified Linear Unit (Leaky ReLU) activation function was used throughout the entire network and also after the first batch-normalization layer in the residual blocks. After being passed through the initial layers and residual blocks, the input was transformed by one more convolution and subsequent Leaky ReLU operation. The result of this process was upsampled conventionally using nearest neighbor interpolation with a factor of two. As one would expect from nearest neighbor interpolation, this step creates a 3D volume with the correct output dimensions but the desired sharpness and increase in detail were not present yet. A convolutional layer with a kernel shape of 3 × 3 × 3 was used to attain these qualities in the final output.

#### 5.3.2. Discriminator architecture

The discriminator (Suppl. Fig. 2) was the 3-dimensional equivalent to a typical convolutional network used for binary image classification. It comprises eight 3D convolution layers, after seven of which batch normalization and ReLU were utilized. After the very first 3D convolution, no batch normalization is performed before applying the activation function. Strided convolution was used as a more dynamic alternative to pooling^50^. Just like in the generator, the kernels had a shape of 3 × 3 × 3 and Leaky ReLU was used as the activation function. Starting at 32, the number of output feature-maps was doubled with every convolutional layer. Finally, two fully connected layers followed by a sigmoid activation layer were used to predict the probability of the provided image being either real or fake. While the generator had a purely convolutional and hence size invariant network structure, the discriminator relies on fully connected layers, causing this part of the adversarial network to be applicable solely to volumes with the same dimensions as those in the training dataset.

### 5.4. Loss function

The loss function of the generator was composed of three terms, the first two of which were similarity measures between the voxel intensities in the original image *I_HR_* and its upsampled pair *I_SR_*,

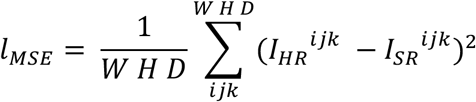

And

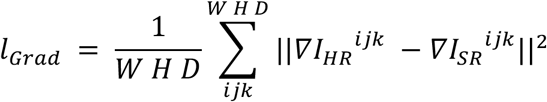

where W, H, D denotes the width, height, and depth of the MR image, *ijk* indicates the voxel index, and the del operator ∇ indicates the spatial gradient of the image. Additionally, the adversarial component of the loss was defined as

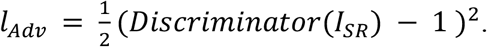

The total loss for the generator is then given by

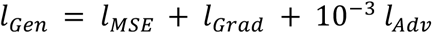

and the discriminator loss is given by

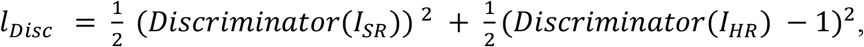

encoding the expected outputs of the *Discriminator*({*I_SR_, I_HR_*}) = {0,1}^51^. GANs can be difficult to train^52^. To aid in their stabilization, data was intentionally mislabeled with 10% probability, and the discriminator output was randomly smoothed^27^. Optimization was performed using one instance of the Adam optimizer^53^ for each of the two networks, as recommended by the TensorFlow documentation. The SRGANs were implemented using TensorFlow 2.4^34^. Training and evaluation were performed on Nvidia P100s GPUs. To accommodate for the impact of random weight initialization, we ran four instances of each training with different initial weights.

### 5.5. Metrics

In order to gauge the image quality of the resulting super sampled images, the peak signal-to-noise ratio (PSNR) and the structural self-similarity index measure (SSIM)^54^ were used. These two metrics have been extensively used both in computer vision and in MR imaging^55^ literature. Scikit-image version 0.18.0^56^ was used to calculate the metric scores. To remove biases due to different brain sizes, these quantities were calculated and averaged only over the unmasked parts of the image. As such, it is difficult to compare the PSNR/SSIM values in the current work with those in most previous works, which have not excluded the background from this calculation.

Although the usage of synthetic metrics is common practice in determining an image’s quality, recently their deployment in medical imaging has come under scrutiny^57^, as they have some prominent issues. For instance, scrambled images can reach high values of SSIM, and PSNR can poorly predict the quality of images as perceived by humans when comparing across different imaging methods^57^. When developing SRGANs for neuroscientific applications, domain-specific metrics closer to real-life applications are also of interest. Including these more neuroanatomical markers allows providing face validity within the neuroimaging domain. Brain volume, white matter surface area, and cortical thickness are just three examples of such complex metrics with scientific relevance. We investigated whether SRGAN-based upsampling interferes with these anatomical metrics for the 1000BRAINS dataset, as well as how different sample sizes affect them. Cortical thickness, surface area, and brain volume were estimated from MR images using FastSurfer^32^. A summary of FastSurfer’s pipeline is provided here: (1) The brain image is segmented into cerebrospinal fluids, gray matter, and white matter. The image is then (2) motion-corrected, (3) intensity normalized and (4) transformed into Talairach space. (5) Tessellation of the gray matter-white matter interface is then performed using the marching cubes algorithm. (6) Topological defects created by the tesselation are fixed. (7) This surface is then expanded until its vertices meet the pial surface. (8) Cortical thickness for a given vertex of the pial surface is measured as the minimal distance to any vertex in the gray matter-white matter interface surface. (9) Surface area is calculated as the sum of the area of each triangle in the tessellation. (10) Brain volume is calculated as the sum of the volumes of all voxels that are not part of the background, ventricles (lateral, inferior lateral, 3rd, 4th, 5th), CSF, or choroid plexus. Two-sided Wilcox t-tests were used for null hypothesis testing.

### 5.6. Sample size scaling

To determine the impact of different dataset sizes on the upsampling results, we compare networks trained on subsets of the 1000BRAINS cohort with sizes 99 (comprising 10% of its training dataset), 247 (25%), 492 (50%), and 985 (100%) subjects. The same evaluation dataset was used for all models and subset sizes. Four different subsets were randomly sampled for each size, except for the dataset containing the whole training data, resulting in a total of 16 training sessions per size. While the number of scans varied, the number of iterations in each training session was kept constant for all training sessions in order to ensure that differences in performance were not caused by training length.

## 6. Conclusions

This work showed that SRGANs can generate high quality upsampled MR images even in the presence of complex, real-life conditions. It also demonstrated that these networks are resilient against population-based artifacts, giving further evidence of their applicability in clinical settings. Considering domain-specific beyond standard synthetic metrics seems to be a relevant step for evaluating the performance of upsampling algorithms on their way towards real-life applications in diagnostics and research.

## Supporting information

Supplementary materials with 3 tables and 2 figures.

## 7. Acknowledgments

The authors gratefully acknowledge the computing time granted through JARA on the supercomputer JURECA^58^ at Forschungszentrum Jülich, which was used for earlier testing. The authors thank Karl-Josef Langen and his team for providing MRI data of brain tumor patients. This project has received funding from the European Union’s Horizon 2020 Research and Innovation Programme under Grant Agreement No. 945539 (HBP SGA3; SC). This research was supported by the Joint Lab “Supercomputing and Modeling for the Human Brain”.

## 8. Data availability

1000BRAINS data is available via applications as specified in the design paper^45^. dHCP data is available upon registration from http://www.developingconnectome.org/data-release. The brain tumor data is available upon reasonable request from the responsible PI (MK).

## 9. Code availability

The code for all experiments is available at <AVAILABLE UPON ACCEPTANCE>. The code for *brainhance* is available at https://github.com/LucasCampos/brainhance^33^.

## References

1. Plenge, E. et al. Super-resolution methods in MRI: Can they improve the trade-off between resolution, signal-to-noise ratio, and acquisition time? Magn. Reson. Med. 68, 1983–1993 (2012).

2. Wolbarst, A. B. & Yanasak, N. An introduction to MRI for medical physicists and engineers. (2019).

3. Lehmann, T. M., Gonner, C. & Spitzer, K. Survey: interpolation methods in medical image processing. IEEE Trans. Med. Imaging 18, 1049–1075 (1999).

4. Jog, A., Carass, A. & Prince, J. L. Self Super-Resolution for Magnetic Resonance Images. in Medical Image Computing and Computer-Assisted Intervention - MICCAI 2016 (eds. Ourselin, S., Joskowicz, L., Sabuncu, M. R., Unal, G. & Wells, W.) vol. 9902 553–560 (Springer International Publishing, 2016).

5. Manjón, J. V. et al. Non-local MRI upsampling. Med. Image Anal. 14, 784–792 (2010).

6. Rousseau, F. Brain Hallucination. in Computer Vision – ECCV 2008 (eds. Forsyth, D., Torr, P. & Zisserman, A.) vol. 5302 497–508 (Springer Berlin Heidelberg, 2008).

7. LeCun, Y., Bengio, Y. & Hinton, G. Deep learning. Nature 521, 436–444 (2015).

8. Krizhevsky, A., Sutskever, I. & Hinton, G. E. ImageNet classification with deep convolutional neural networks. Commun. ACM 60, 84–90 (2017).

9. Matek, C., Schwarz, S., Spiekermann, K. & Marr, C. Human-level recognition of blast cells in acute myeloid leukaemia with convolutional neural networks. Nat. Mach. Intell. 1, 538–544 (2019).

10. Ronneberger, O., Fischer, P. & Brox, T. U-Net: Convolutional Networks for Biomedical Image Segmentation. ArXiv150504597 Cs (2015).

11. Kingma, D. P. & Welling, M. Auto-Encoding Variational Bayes. ArXiv13126114 Cs Stat (2014).

12. Wang, X., Wang, K. & Lian, S. A survey on face data augmentation for the training of deep neural networks. Neural Comput. Appl. 32, 15503–15531 (2020).

13. Xu, W., Keshmiri, S. & Wang, G. Adversarially Approximated Autoencoder for Image Generation and Manipulation. IEEE Trans. Multimed. 21, 2387–2396 (2019).

14. Dong, C., Loy, C. C., He, K. & Tang, X. Learning a Deep Convolutional Network for Image Super-Resolution. in Computer Vision –ECCV 2014 (eds. Fleet, D., Pajdla, T., Schiele, B. & Tuytelaars, T.) vol. 8692 184–199 (Springer International Publishing, 2014).

15. Dong, C., Loy, C. C., He, K. & Tang, X. Image Super-Resolution Using Deep Convolutional Networks. IEEE Trans. Pattern Anal. Mach. Intell. 38, 295–307 (2016).

16. Pham, C.-H., Ducournau, A., Fablet, R. & Rousseau, F. Brain MRI super-resolution using deep 3D convolutional networks. in 2017 IEEE 14th International Symposium on Biomedical Imaging (ISBI 2017) 197–200 (IEEE, 2017). doi:10.1109/ISBI.2017.7950500.

17. Du, J. et al. Super-resolution reconstruction of single anisotropic 3D MR images using residual convolutional neural network. NEUROCOMPUTING 392, 209–220 (2020).

18. Du, J. et al. Brain MRI super-resolution using 3D dilated convolutional encoder-decoder network. IEEE ACCESS 8, 18938–18950 (2020).

19. Jurek, J., Kocinski, M., Materka, A., Elgalal, M. & Majos, A. CNN-based superresolution reconstruction of 3D MR images using thick-slice scans. Biocybern. Biomed. Eng. 40, 111–125 (2020).

20. Pham, C.-H. et al. Multiscale brain MRI super-resolution using deep 3D convolutional networks. Comput. Med. IMAGING Graph. 77, (2019).

21. Goodfellow, I. et al. Generative adversarial nets. in Advances in neural information processing systems 27 (eds. Ghahramani, Z., Welling, M., Cortes, C., Lawrence, N. D. & Weinberger, K. Q.) 2672–2680 (Curran Associates, Inc., 2014).

22. Yi, X., Walia, E. & Babyn, P. Generative adversarial network in medical imaging: A review. Med. Image Anal. 58, 101552 (2019).

23. Yang, W. et al. Deep Learning for Single Image Super-Resolution: A Brief Review. IEEE Trans. Multimed. 21, 3106–3121 (2019).

24. Abramian, D. & Eklund, A. Refacing: Reconstructing Anonymized Facial Features Using GANS. in 2019 IEEE 16th International Symposium on Biomedical Imaging (ISBI 2019) 1104–1108 (IEEE, 2019). doi:10.1109/ISBI.2019.8759515.

25. Dar, S. UH. et al. Image Synthesis in Multi-Contrast MRI With Conditional Generative Adversarial Networks. IEEE Trans. Med. Imaging 38, 2375–2388 (2019).

26. Nie, D. et al. Medical Image Synthesis with Context-Aware Generative Adversarial Networks. in Medical Image Computing and Computer Assisted Intervention – MICCAI 2017 (eds. Descoteaux, M. et al.) vol. 10435 417–425 (Springer International Publishing, 2017).

27. Sánchez, I. & Vilaplana, V. Brain MRI super-resolution using 3D generative adversarial networks. in (2018).

28. Chen, Y. et al. Brain MRI super resolution using 3D deep densely connected neural networks. in 2018 IEEE 15th International Symposium on Biomedical Imaging (ISBI 2018) 739–742 (IEEE, 2018).

29. Isola, P., Zhu, J.-Y., Zhou, T. & Efros, A. A. Image-to-Image Translation with Conditional Adversarial Networks. ArXiv161107004 Cs (2018).

30. Ledig, C. et al. Photo-Realistic Single Image Super-Resolution Using a Generative Adversarial Network. ArXiv160904802 Cs Stat (2017).

31. Cohen, J. P., Luck, M. & Honari, S. Distribution Matching Losses Can Hallucinate Features in Medical Image Translation. in Medical Image Computing and Computer Assisted Intervention – MICCAI 2018 (eds. Frangi, A. F., Schnabel, J. A., Davatzikos, C., Alberola-López, C. & Fichtinger, G.) 529–536 (Springer International Publishing, 2018). doi:10.1007/978-3-030-00928-1_60.

32. Henschel, L. et al. FastSurfer - A fast and accurate deep learning based neuroimaging pipeline. NeuroImage 219, 117012 (2020).

33. Campos, L. & Förster, L. LucasCampos/brainhance: v0.0.3. (Zenodo, 2022). doi:10.5281/ZENODO.6624433.

34. Abadi, M. et al. TensorFlow: Large-scale machine learning on heterogeneous systems. (2015).

35. Cherukuri, V., Guo, T., Schiff, S. J. & Monga, V. Deep MR brain image super-resolution using spatio-structural priors. IEEE Trans. IMAGE Process. 29, 1368–1383 (2020).

36. Du, J., Wang, L., Gholipour, A., He, Z. & Jia, Y. Accelerated super-resolution MR image reconstruction via a 3D densely connected deep convolutional neural network. in PROCEEDINGS 2018 IEEE INTERNATIONAL CONFERENCE ON BIOINFORMATICS AND BIOMEDICINE (BIBM) (ed. Zheng, H and Callejas, Z and Griol, D and Wang, H and Hu, X and Schmidt, H and Baumbach, J and Dickerson, J and Zhang, L) 349–355 (2018).

37. Park, J. et al. Computed tomography super-resolution using deep convolutional neural network. Phys. Med. Biol. 63, 145011 (2018).

38. Shi, J. et al. MR image super-resolution via wide residual networks with fixed skip connection. IEEE J. Biomed. Health Inform. 23, 1129–1140 (2019).

39. Zeng, K. et al. Simultaneous single-and multi-contrast super-resolution for brain MRI images based on a convolutional neural network. Comput. Biol. Med. 99, 133–141 (2018).

40. Delannoy, Q. et al. SegSRGAN: Super-resolution and segmentation using generative adversarial networks - Application to neonatal brain MRI. Comput. Biol. Med. 120, (2020).

41. Jahng, G.-H., Li, K.-L., Ostergaard, L. & Calamante, F. Perfusion Magnetic Resonance Imaging: A Comprehensive Update on Principles and Techniques. Korean J. Radiol. 15, 554 (2014).

42. Shi, W. et al. Cardiac Image Super-Resolution with Global Correspondence Using Multi-Atlas PatchMatch. in Medical Image Computing and Computer-Assisted Intervention – MICCAI 2013 (eds. Mori, K., Sakuma, I., Sato, Y., Barillot, C. & Navab, N.) vol. 8151 9–16 (Springer Berlin Heidelberg, 2013).

43. Kocher, M. et al. Role of the default mode resting-state network for cognitive functioning in malignant glioma patients following multimodal treatment. NeuroImage Clin. 27, 102287 (2020).

44. Makropoulos, A. et al. The developing human connectome project: A minimal processing pipeline for neonatal cortical surface reconstruction. NeuroImage 173, 88–112 (2018).

45. Caspers, S. et al. Studying variability in human brain aging in a population-based German cohort—rationale and design of 1000BRAINS. Front. Aging Neurosci. 6, (2014).

46. Garyfallidis, E. et al. Dipy, a library for the analysis of diffusion MRI data. Front. Neuroinformatics 8, (2014).

47. Isensee, F. et al. Automated brain extraction of multisequence MRI using artificial neural networks. Hum. Brain Mapp. 40, 4952–4964 (2019).

48. Tournier, J.-D. et al. MRtrix3: A fast, flexible and open software framework for medical image processing and visualisation. NeuroImage 202, 116137 (2019).

49. Virtanen, P. et al. SciPy 1.0: Fundamental algorithms for scientific computing in python. Nat. Methods 17, 261–272 (2020).

50. Springenberg, J. T., Dosovitskiy, A., Brox, T. & Riedmiller, M. Striving for Simplicity: The All Convolutional Net. ArXiv14126806 Cs (2015).

51. Mao, X. et al. Least Squares Generative Adversarial Networks. (2017).

52. Thanh-Tung, H., Tran, T. & Venkatesh, S. Improving Generalization and Stability of Generative Adversarial Networks. ArXiv190203984 Cs Stat (2019).

53. Kingma, D. P. & Ba, J. Adam: A Method for Stochastic Optimization. ArXiv14126980 Cs (2017).

54. Wang, Z., Bovik, A. C., Sheikh, H. R. & Simoncelli, E. P. Image Quality Assessment: From Error Visibility to Structural Similarity. IEEE Trans. Image Process. 13, 600–612 (2004).

55. Kumar, B., Kumar, S. B. & Kumar, C. Development of improved SSIM quality index for compressed medical images. in 2013 IEEE Second International Conference on Image Information Processing (ICIIP-2013) 251–255 (IEEE, 2013). doi:10.1109/ICIIP.2013.6707593.

56. van der Walt, S. et al. scikit-image: image processing in Python. PeerJ 2, e453 (2014).

57. Pambrun, J.-F. & Noumeir, R. Limitations of the SSIM quality metric in the context of diagnostic imaging. in 2015 IEEE International Conference on Image Processing (ICIP) 2960–2963 (IEEE, 2015). doi:10.1109/ICIP.2015.7351345.

58. Thörnig, P. JURECA: Data Centric and Booster Modules implementing the Modular Supercomputing Architecture at Jülich Supercomputing Centre. J. Large-Scale Res. Facil. JLSRF 7, A182 (2021).

