## Supplementary materials with 3 tables and 2 figures. for "Towards robust and generalizable super-resolution generative adversarial networks for magnetic resonance neuroimaging: a cross-population approach"

### Supplementary Information

#### S1 Cohort sizes

Table S1: Size of cohorts. The cohorts were split into a training dataset comprising approximately 80% of the data and an evaluation dataset comprising the remaining 20%.

| **Dataset** | **Total** | **Training** | **Evaluation** |
|---|---|---|---|
| 1000BRAINS | 1233 | 986 | 247 |
| Brain Tumor | 187 | 149 | 38 |
| dHCP | 451 | 360 | 91 |
| Combined | 1871 | 1495 | 376 |

**S2 Hyperparameters**

Table S2: Hyperparameters used during training

| **Parameter** | **Value** |
|---|---|
| Optimizer | Adam |
| Learning rate discriminator | $10^{-6}$ |
| Learning rate discriminator | $10^{-4}$ |
| Learning rate steps | 4920 |
| Learning decay | 0.99 |
| Batch size | 2 |

**S3 Anatomical measurements with different sample size**

Table S3: Comparison of anatomical measurements for training sessions comprising increasing number of subjects (Experiment 1). Asterisks indicate the level of statistical significance (ns: $p > 0.05$, $*$ : $p \leq 0.05$, $**$ : $p \leq 0.01$, $***$ : $p \leq 0.001$). All values are given as mean ± std. n = 247.

| | **Diff to HighRes** |
|---|---|
| **SRGANs (99 subjects)** | |
| Cortical Thick. (LH) | -18.20±1.92 *** |
| Cortical Thick. (RH) | -17.38±2.33 *** |
| Cortical Area (LH) | 2.52±4.55 *** |
| Cortical Area (RH) | -1.12±5.08 *** |
| Brain Volume | 0.09±5.69 ns |
| **SRGANs (247 subjects)** | |
| Cortical Thick. (LH) | -18.75±2.16 *** |
| Cortical Thick. (RH) | -18.26±2.43 *** |
| Cortical Area (LH) | 2.04±5.47 *** |
| Cortical Area (RH) | -1.21±6.18 *** |
| Brain Volume | 0.06±6.07 ns |
| **SRGANs (492 subjects)** | |
| Cortical Thick. (LH) | -19.84±2.12 *** |
| Cortical Thick. (RH) | -19.03±2.37 ns |
| Cortical Area (LH) | 2.43±4.91 *** |
| Cortical Area (RH) | -0.93±5.54 ** |
| Brain Volume | 0.13±6.19 * |
| **SRGANs (985 subjects)** | |
| Cortical Thick. (LH) | -18.94±2.01 *** |
| Cortical Thick. (RH) | -18.18±2.33 *** |
| Cortical Area (LH) | 2.29±3.71 *** |
| Cortical Area (RH) | -0.65±3.95 ** |
| Brain Volume | 0.95±5.17 * |

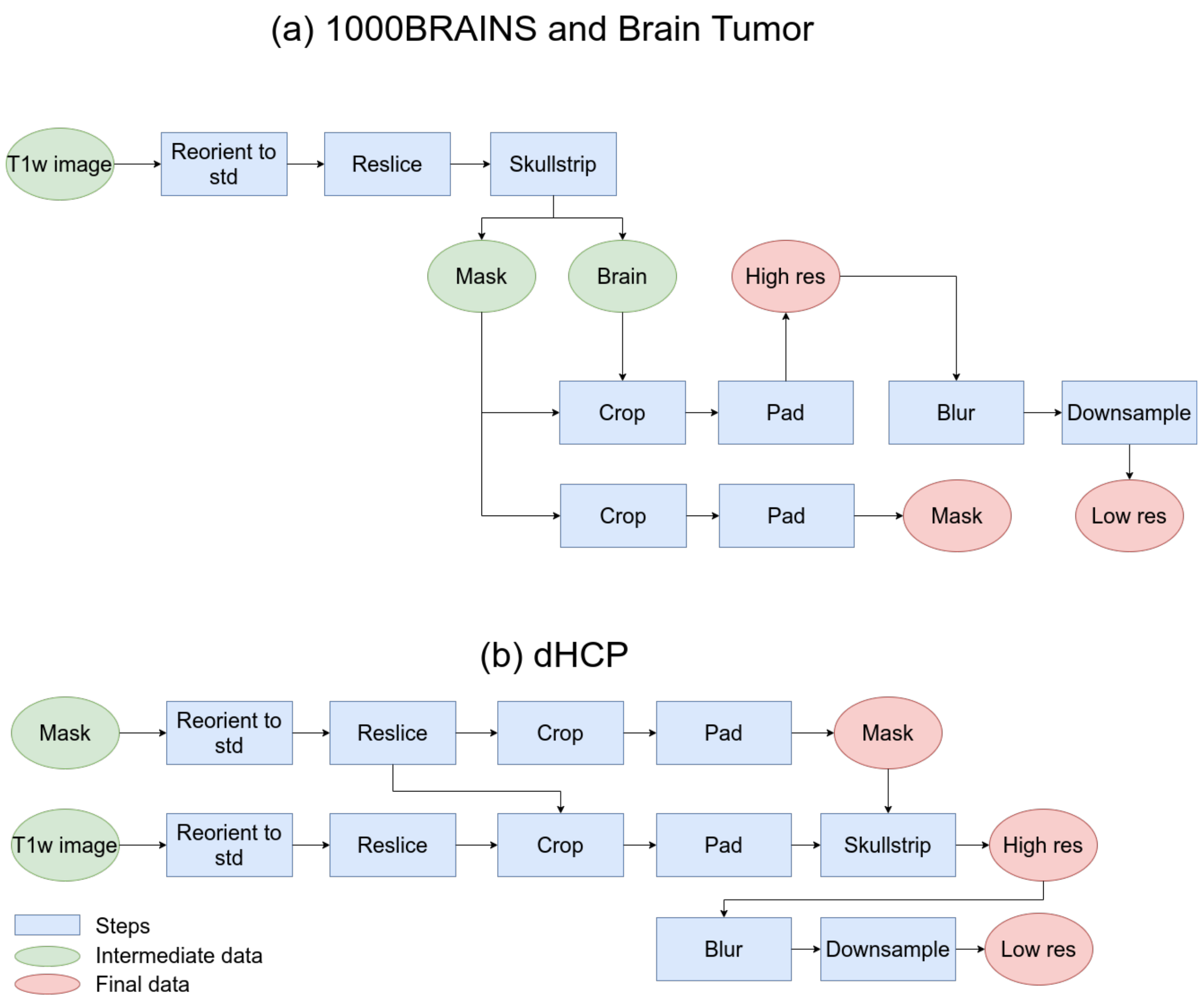


Supplementary Figure 1: Preprocessing pipeline for (a) the 1000BRAINS and Brain Tumor cohorts and (b) the dHCP cohort. [Full width]

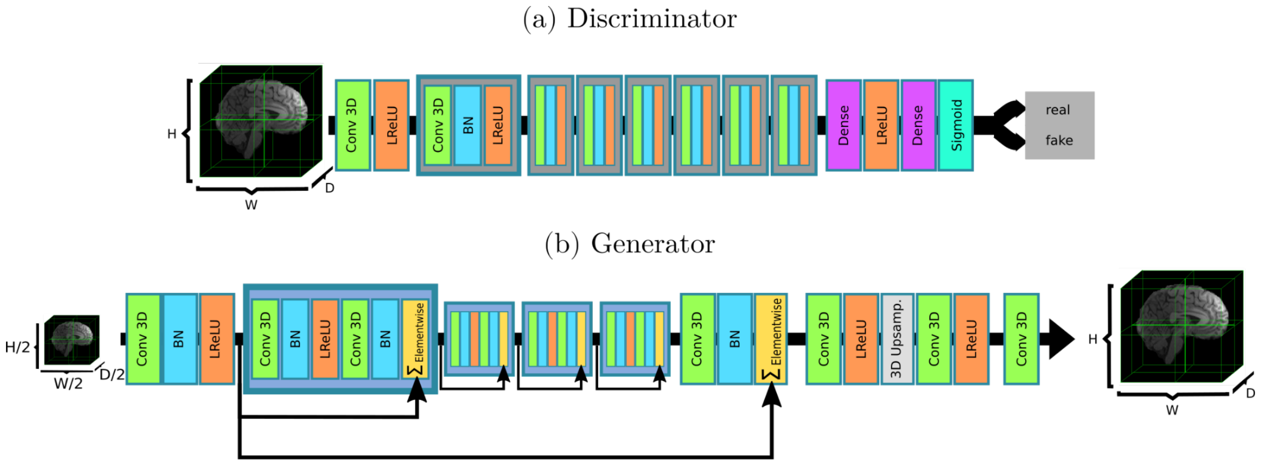


Supplementary Figure 2: Network architecture for (a) discriminator, and (b) generator networks. Based on Ref.[1]. [Full width]

### References

1. Ledig, C. *et al.* Photo-Realistic Single Image Super-Resolution Using a Generative Adversarial Network. *ArXiv160904802 Cs Stat* (2017).
